# Parametric effects of light acting via multiple photoreceptors contribute to circadian entrainment in *Drosophila melanogaster*

**DOI:** 10.1101/2022.03.02.482722

**Authors:** Lakshman Abhilash, Orie Thomas Shafer

**Affiliations:** The Advanced Science Research Center, The City University of New York; The Graduate Center at the City University of New York

**Keywords:** parametric entrainment, continuous entrainment, tonic entrainment, Drosophila, circadian, locomotor activity, Aschoff’s rule, free-running period, constant light, skeleton photoperiods

## Abstract

Circadian rhythms in physiology and behavior have near 24-hour periodicities that must adjust to the exact 24-hour geophysical cycles on earth to ensure adaptive daily timing. Such adjustment is called entrainment. One major mode of entrainment is via the continuous modulation of circadian period by the prolonged presence of light. Although *Drosophila melanogaster* is a prominent insect model of chronobiology, there is little evidence for such continuous effects of light in the species. In this study, we demonstrate that prolonged light exposure at specific times of the day shapes the daily timing of activity in flies. We also establish that continuous blue- and UV-blocked light lengthens the circadian period of *Drosophila* and provide evidence that this is produced by the combined action of multiple photoreceptors which, includes the cell autonomous photoreceptor *cryptochrome*. Finally, we introduce ramped light cycles as an entrainment paradigm that produces light entrainment that lacks the large light-driven startle responses typically displayed by flies and requires multiple days for entrainment to shifted cycles. These features are reminiscent of entrainment in mammalian models systems and make possible new experimental approaches to understanding the mechanisms underlying entrainment in the fly.

## Introduction

Circadian rhythms are ubiquitous endogenous rhythms in behavior, physiology and metabolism. Such rhythms “free-run” in the absence of cyclic environments at periodicities close to but significantly different from 24-h. To support optimal daily timing on a rotating planet, these rhythms must be coordinated with the exact 24-h rhythms of the earth’s geophysical cycles, which include daily changes in light and temperature. Such coordination, called entrainment, enables organisms to restrict their daily activities and other biological processes to appropriate times of the day and to anticipate predictable daily changes in the environment [1–4]. Poor entrainment contributes significantly to many health problems in modern societies, making entrainment a central problem to circadian biology and general well-being [4].

Organisms are thought to entrain to cyclic environments using two distinct processes [1,5]. In the first, referred to as non-parametric, environmental transitions instantaneously reset the phase of the circadian clock, much like watches being reset upon the switch to daylight saving time [1,6]. In the second, referred to as parametric, the continuous action of light serves to speed-up or slow-down the angular velocity of internal clocks in a time-of-day dependent manner [1,5,7]. Real world entrainment likely involves both parametric and non-parametric processes [5] and understanding both processes is critical if we are to understand the entrainment of the circadian system and how clocks operate under both natural and artificial light environments.

Both parametric and non-parametric effects of light have been incorporated into unified models of circadian entrainment, including for *Drosophila*, an organism that offers a powerful model system for understanding the neuronal and molecular basis of light entrainment [8–10]. However, the fly’s circadian system does not maintain robust endogenous circadian rhythms under constant light (LL), making the mechanisms of parametric entrainment difficult to address. This is because of light and CRYPTOCHROME (CRY; blue light photoreceptor) mediated degradation of TIMELESS (TIM), a core component of the molecular circadian clock ([11], reviewed in [12]). Indeed, very few studies have examined the parametric effects of light on the fly clock. Free-running rhythms have been observed for *Drosophila*, both in *cryptochrome* loss of function mutations, which maintain rhythms in LL [13–16] and in small numbers of wild-type flies [17,18]. These studies suggest that LL conditions lengthen the clock’s endogenous period relative to constant darkness (DD). However, the sensitivity of the fly clock to light means that LL conditions most often cause arrhythmicity or internal desynchronization in wild-type strains [15] and this has limited the utility of the fly as a model of parametric light entrainment. For this reason, we have sought to develop approaches that allow us to use the powerful fly model to understand the parametric effects of light.

We therefore set out to identify light regimes that better support an examination of parametric light effects and their contribution to circadian entrainment in *Drosophila*. Using various photoperiodic regimes (see Methods) we show that the timing of locomotor activity under standard laboratory conditions is strongly shaped by long durations of light at specific times of the day. This suggested that speed of the clock regulating these rhythms must be modulated by time-of-day dependent continuous effects of light. In our search for LL conditions that support free-running rhythms in the fly, we find that LL regimes in which UV and blue wavelengths have been blocked support robust free-running rhythms and produce lengthened circadian periods (slower running clocks).

We also show that external photoreceptors and the blue light circadian photoreceptor *cryptochrome* differentially govern the parametric effects of light under these conditions. Finally, we define light regimes that entrain behavioral rhythms in a manner that fails to induce the confounding startle responses associated with widely used LD12:12 cycles (12-h of light and 12-h of darkness) and that, as in mammalian systems, such regimes require multiple cycles for entrainment to shifted light cycles. These results reveal important features of entrainment in *Drosophila* and establish a useful new approach to understanding light resetting in this important model organism.

## Methods

### Fly stocks

The following fly lines were used in this study: *Canton-S* (*CS*; Bloomington Drosophila Stock Center (BDSC) stock number: 64349), white-eye (*w^1118^*) [19], yellow-white (*yw*; BDSC stock number: 1495), *yw;;cry^OUT^* [20], *w;;glass^[60j]^* [13], *+;;norpA^[7]^* (BDSC stock number: 5685), *+;;per^01^* (BDSC stock number: 80928) and *+;;clk^JRK^* (BDSC stock number: 80927). All these genotypes were maintained in bottles with corn syrup soy media (Archon Scientific; Durham, NC). Three to five-day old male flies were collected from bottles for loading into *Drosophila* Activity Monitors (DAM; Trikinetics, Waltham, MA). Sample sizes and replication for each experiment and the environmental conditions experienced before behavioral experiments began are described separately for each experiment below.

### Entrainment of locomotor activity rhythms under skeleton photoperiods

#### Fly lines, recording conditions and light conditions

*CS*, *w^1118^* and *yw* males were used in these experiments. Flies were maintained under white light LD12:12 before being loaded into locomotor tubes for one cycle after being loaded into the DAM systems before being transferred to the different skeleton photoperiod conditions. ~32 flies of each genotype were collected under CO_2_ anesthesia and gently placed into 5-mm glass capillary tubes containing sucrose-agar media, which was sealed with paraffin wax at the media containing end and plugged with a short length of white yarn. Fly-containing tubes were then loaded into DAM monitors and secured with rubber bands woven between tubes. Infrared beam crossings were recorded every minute for the duration of the experiment. Our symmetric skeleton photoperiod (SPP; Fig. 1A), consisted of 30-min pulses starting and ending at dawn (Zeitgeber Time (ZT) 00) and dusk (ZT 12) of the LD cycle used during rearing, respectively. Additionally, we used two asymmetric skeleton photoperiods (aSPP), the first of which consisted of a dawn light pulse lasting 6-h with a 30-min pulse ending at dusk (aSPP-1; Fig. 1A), and the second of which consisted of a dusk light pulse lasting 6-h and a 30-min pulse starting at dawn (aSSP-2; Fig. 1A). All these conditions employed the standard broad spectrum white light produced by the incubators without the use of any wavelength filters. These incubators were programmed to run at a constant 25 °C. The light intensity for all these experiments was 400-500 lux.

**Figure 1:**
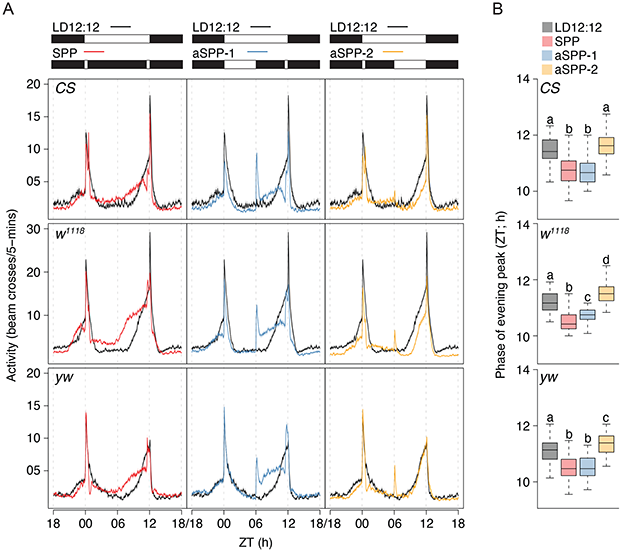
Entrainment to white light skeleton photoperiods. (A) Mean (±SEM) locomotor activity profiles of *CS* (top), *w^1118^* (middle) and *yw* (bottom) flies under symmetric skeleton photoperiods (left) and two asymmetric skeleton photoperiods (middle and right). The photoperiodic regimes are shown on top of the profile plots. Black shaded regions indicate darkness and white regions indicate light phases (400-500 lux) of the 24-h cycle. Note that the profiles under LD12:12 are replotted across panels to facilitate pair-wise comparisons. (B) Phases of the evening peak of activity for the three strains across the different photoperiodic conditions are shown. Boxplots that share the same letter are not statistically significantly different from each other. Statistical comparison of phases across all four photoperiodic conditions were done using Kruskal-Wallis tests. *CS*: 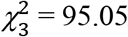, *p* = 0; 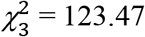, *p* = 0; *yw*: 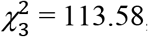, *p* = 0. Profiles and phases of evening peak of activity are pooled from two independent replicate runs for each genotype.

#### Statistical analyses

The first step in the analysis of these data was to identify flies that were entrained versus flies that were free-running. Owing to the strong startle responses associated with step-function cycles, especially under the skeleton photoperiod conditions, it was not possible to use raw data to perform periodogram analyses and estimate period values. This is because startles to transitions (see Fig. 1A) cause a spike in activity at recurring 24-h intervals which produce strong 24-h signals in standard periodogram analyses, even when the non-startled component of activity appear to be free-running. Therefore, we used the Savitzky-Golay (SG) filter with a filter order of 2 and filter frame length of 251-min to smooth the raw data, to minimize the contributions of startle to the periodogram results. We then used the *χ^2^* periodogram on the filtered data and estimated period values. Flies that had period values between 23.67-h and 24.33-h were considered entrained, and the ones outside this range were considered free-running. These values were chosen because they’re the two closest points to 24-h (on either side) in our *χ^2^* periodogram analysis; this enabled us to include flies that entrain but have periodicities that may be slightly wavering around 24-h.

Profiles of activity were averaged across only the flies that were categorized as entrained. Phases of evening peaks of activity were estimated by applying the SG filter on raw data of entrained flies only. For this purpose, the filter order was kept at two, but the filter frame length was reduced to 81-min so that small nuances of peaks could be identified. For each genotype, the effect of light environment on phases of evening peaks of activity were assessed by Kruskal-Wallis tests followed by Bonferroni corrections to account for multiple comparisons from the *agricolae* package for R [21]. All the analyses were carried out using custom R scripts.

### Assessment of free-running period under constant light

#### Fly lines, recording conditions and the constant light paradigm

*CS*, *w^1118^*, *yw*, *cry^OUT^*, *glass^[60j]^* and *norpA^[7]^* males were used in these experiments. ~32 flies of each genotype were collected and loaded into locomotor tubes as described above. Infrared beam crossings were recorded every minute for at least 10 days under constant darkness (DD) or constant light (LL) in Percival incubators (Model: LED-30HL1, Geneva Scientific, Fontana, WI) at 25 °C. Data from at least two independent runs are reported. See Suppl. Table 1 for sample sizes and number of replicate runs.

Given that wildtype *Drosophila* are arrhythmic under constant white light, we sought to establish light conditions under which wild-type flies maintain robust free-running rhythms. It is well established that the arrhythmicity of flies under white LL is driven by the Cryptochrome (CRY) mediated TIMELESS (TIM) degradation [22,23]. Furthermore, CRY is maximally sensitive to blue and UV wavelengths of light [24]. We therefore used multiple light filters which had varying degrees of overlap with the CRY action spectrum (Fig. 2A). We used constant light of green (Filter number: 139; 8-12 umol/m2/sec), orange (Filter number: 105), true yellow (Filter number: 101; 8-12 umol/m2/sec) and blue and UV-blocked (Filter number: 821) wavelengths by means of using specific filters (Lee Filters, UK; http://www.leefilters.com/). We also used constant red light using the in-built red LED in our Percival incubators (Fig. 2A). In all cases, we used light intensities between 6-12 *μmol/m^2^/sec*. For examining the dose dependent effect of blue and UV-blocked light we used three intensities, i.e., 14-18 (~16) *μW/cm^2^*, 28-32 (~30) *μW/cm^2^* and 47-53 (~50) *μW/cm^2^* at 600nm. We used the S120C sensor and the PM100D console from Thorlabs (Newton, NJ; https://www.thorlabs.com/) to measure the single wavelength light intensities. Comparisons of period for various genotypes reported in Figs. 3B and 3C (and Suppl. Fig. 1) under blue and UV-blocked light were done after assaying flies under intensities of 30-45 *μW/cm^2^* at 600nm.

**Figure 2:**
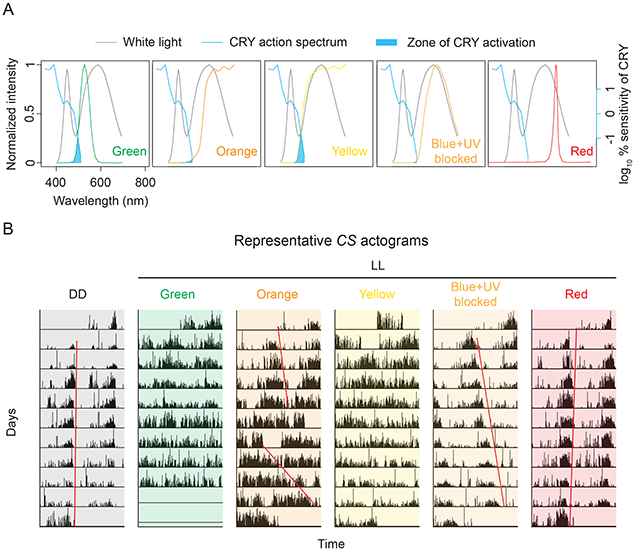
Locomotor activity rhythms under various constant light conditions. (A) Spectral composition of broad-spectrum white light overlayed with spectral composition of lights when various filters were used, and with the action spectrum of CRYPTOCHROME (CRY). Data for the spectral composition were extracted from the Lee Filters website (see Methods) and for the action spectrum of CRY were taken from VanVickle-Chavez and Van Gelder, 2007 [24]. (B) Example actograms of *CS* flies under constant darkness and constant light of various wavelength compositions. Red lines on the actograms are eye-fit lines through the offset of locomotor activity.

**Figure 3:**
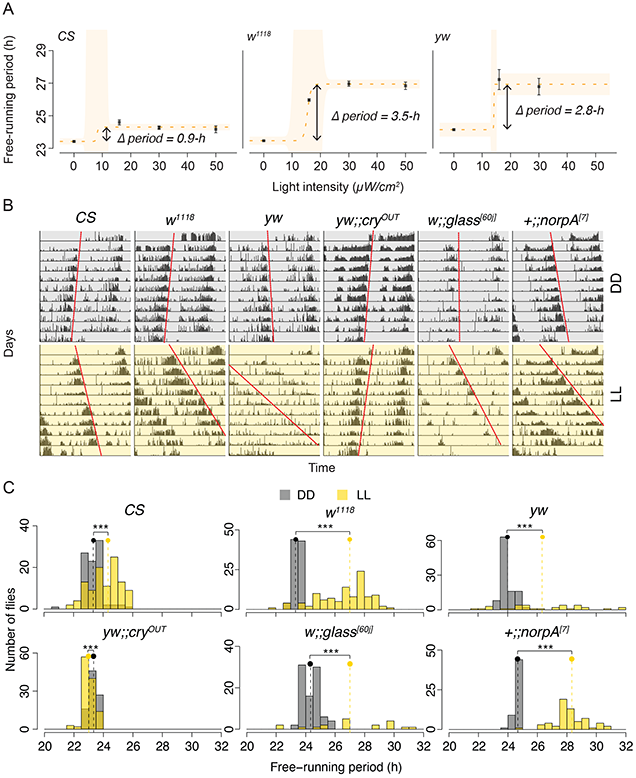
Free-running locomotor activity rhythms of flies under constant blue- and UV-blocked light. (A) Dose-dependent effect of constant light on the free-running period of *CS* (left), *w^1118^* (middle) and *yw* (right) flies. The lack of data point for highest intensity of light in *yw* flies is because flies were arrhythmic under this condition. The yellow dashed line is dose response curve fitted through a logistic model (see Methods). ANOVAs on the fitted logistic model revealed a statistically significant effect of intensity on the free-running period for all three genotypes (*CS*: *p* < 2e-16; *w^1118^*: *p* < 2e-16; *yw*: *p* < 2e-16). (B) Representative actograms of *Canton-S* (*CS*), *w^1118^*, *yw*, *yw;;cry^OUT^*, *w;;glass^[60j]^* and *+;;norpA^[7]^* flies under constant darkness (top) and constant blue- and UV-blocked light (bottom) of 30-45 *μW/cm^2^*. (C) Frequency distributions of free-running periods of all genotypes under constant dark and constant light conditions. The dots and dashed vertical lines represent the median values of free-running periods under each regime. Statistical comparisons are based on Wilcoxon’s tests (*CS*: *W* = 2676.5, *p* = 6.036e-08; *w^1118^*: *W* = 502, *p* = 2.2e-16; *yw*: *W* = 618.5, *p* = 1.583e-08; *yw;;cry^OUT^*: *W* = 7391, *p* = 2.233e-08; *w;;glass^[60j]^*: *W* = 305, *p* = 1.092e-05; *+;;norpA^[7]^*: *W* = 10, *p* = 2.2e-16). Values are pooled from at least two independent runs (see Suppl. Table 1).

#### Statistical analyses

Free-running period values and corresponding rhythmic power were estimated using the *χ^2^* periodogram (implemented using in-house custom R scripts). Period and power values were pooled from all independent replicate runs and were compared using Wilcoxon’s tests for two independent samples (also implemented using custom R scripts). Dose response analyses were carried out using functions from the *drda* package [25] on R using logistic models. The logistic model is a form of non-linear (sigmoidal) growth curve that was fit with our observed data using the default four-parameter function. An ANOVA on the fitted model then tests the model against a null hypothesis of a flat line representing the absence of light intensity dependent changes in the free-running period.

### Entrainment to ramped light cycles

#### Fly lines, recording conditions and light conditions

*CS*, *w^1118^*, *yw*, *cry^OUT^*, *glass^[60j]^*, *norpA^[7]^*, *per^01^* and *clk^JRK^* males were used in these experiments. Flies were reared under ramped yellow light cycles in Percival incubators (Model: LED-30HL1, Geneva Scientific, Fontana, WI) at a constant temperature of 25 °C. ~32 males of each genotype were collected for each run and loaded into locomotor tubes as described above. Ramping light cycles were generated such that there was no darkness at any point in the cycle. We programmed gradual light increases from ~10 *μW/cm^2^* to 40-50 *μW/cm^2^* (measured at 600nm) (ZT00-ZT12) followed by gradual light decreases back to ~10 *μW/cm^2^* during the next 12-h to complete the cycle. We settled on these light cycles based on our desire to avoid distinct jumps in light intensity from complete darkness, which produce large startle responses, and to reduce the possibility of any stark transition in light intensity that that could induce non-parametric clock resetting. We recorded beam crossings, as described above, under the ramped light cycles for nine consecutive days, at the end of which flies were kept under constant light starting from the nadir of the ramped cycle (~10 *μW/cm^2^* intensity) to assess phase control. If entrained, flies must have a similar phase of locomotor activity on the first day under constant conditions as they did during entrainment. This phenomenon, referred to as phase-control, is a pre-requisite for a rhythm to be considered entrained [2].

#### Statistical analyses

Owing to the novelty of this light regime, it was necessary to assess if flies could entrain to sit at all. For each fly, we plotted an actogram over nine ramp cycles, computed a wavelet power spectrum to get daily estimates of period and plotted the standard *χ^2^* periodogram. The wavelet power spectrum was computed using the *WaveletComp* package on R [26] and is appropriate for use with circadian time series [27,28]. Individual flies were considered “entrained” only if visual inspection of the actogram showed that timing of activity had stabilized by the sixth ramping cycle, if the wavelet power spectrum showed stable period values between 23.5-24.5-h post the sixth cycle, and/or if the *χ^2^* periodogram showed peaks in the same range. A fly was categorized as “transiently entrained” or “relatively coordinated” if there were obvious period changes during the time-course (for instance, a period that shifted from period of < 24-h to > 24-h) in the actograms, if there was no period stability established by the sixth cycle based on the wavelet power spectrum, if there were multiple peaks in the *χ^2^* periodogram, or if flies appeared entrained for the first few cycles and then appeared to free-run. Flies were deemed “free-running” if the actograms showed activity bouts drifting in a single direction and if the *χ^2^* periodogram showed peaks at period values less than 23.5 or greater than 24.5-h. Flies were considered “arrhythmic” if there was no significant peak in the *χ^2^* periodogram or if no detectable period in the wavelet power spectrum. For loss-of-function clock mutants, flies were considered rhythmic if there was a significant peak in the *χ^2^* periodogram or if there was a detectable period in the wavelet power spectrum. All these analyses were carried out using custom R scripts.

Only the flies categorized as entrained (or rhythmic, in case of loss-of-function clock mutants) were used to construct the average profiles of activity and to assess phase control under constant dim light. We estimated phases of activity peaks using the SG filter for the entrained flies under cyclic conditions and on the first day of constant dim light. We used a filter order of one and filter frame length of 41 on activity data binned in 15-min intervals to smooth the data before identifying peaks. Phase control was assessed by *V*-tests implemented through the *CircStats* package for R [29]. *V*-tests are Rayleigh’s tests of uniformity on circular data that have an alternate hypothesis of a known mean angle [30]. Difference in phases of peaks between *CS* and *per^01^* flies was tested using a Wilcoxon’s test, also implemented on R.

## Results

### Pulses of light at dawn and dusk are insufficient to mirror the entrained waveform of locomotor activity under standard LD12:12 conditions

One major prediction of the non-parametric (a.k.a., instantaneous) model of entrainment is that light presented at dawn and dusk are the most critical signals for entrainment, and that light during the rest of the day is largely superfluous [5,6]. If this were true of *D. melanogaster* locomotor rhythms, we would expect the locomotor activity waveform of flies be highly similar under LD12:12 and symmetric skeleton photoperiods (SPP), in which only two 30-min light pulses are provided to flies in an otherwise dark environment (see Methods; Fig. 1A). Although such experiments have been done on *Drosophila* eclosion rhythms [31,32], ours is the first, to the best of our knowledge, to report results of such experiments on the locomotor activity rhythm in *D. melanogaster*. All wild-type and genetic background control strains we examined readily entrained to SPP displaying no differences from LD12:12 flies in the percentage of entrained flies (Suppl. Table 2).

Averaged locomotor activity profiles of *CS*, *w^1118^* and *yw* flies indicate that there were distinct differences in waveforms between LD12:12 and SPP (Fig. 1A). We quantified the phases of evening activity peaks from smoothed data (see Methods) and found that median phases in all three genotypes were significantly advanced under SPP compared to LD12:12 (See Methods for details of statistical analysis; Fig. 1B). These results clearly show that skeleton photoperiods do not recapitulate entrainment under LD12:12 and that light in the middle of the day has significant effects on the phase of evening activity.

### Long durations of light only during the second half of the day shape the timing of locomotor activity under standard LD cycles

If parametric effects contribute to entrainment under step function LD cycles, it is expected that increased durations of skeleton light exposure will have measurable effects on the phase of entrainment in a manner that depends on the timing of the increased light [7,32]. The parametric model of entrainment posits that long durations of light delivered at dawn will accelerate the clock, whereas light delivered at dusk will have a decelerating effect on the clock [7,33]. The prediction, therefore, is that long durations of light around dawn would lead to reduced free-running period, thereby leading to an advanced phase of entrainment and that long durations of light during dusk would lengthen free-running period and consequently delay the phase of entrainment [34]. In contrast, if non-parametric effects dominate entrainment at transitions, activity profiles under skeleton light cycles should be largely unaffected by the length of the skeleton light periods. In order to test the extent to which the fly is entrained to LD cycles via parametric or non-parametric effects, we subjected our flies to two asymmetric skeleton photoperiods (aSPP), one with 6-h of light during the first half of the day and a 30-min pulse at dusk (30-min; aSPP-1; Fig. 1A), and one with a 30-min dawn pulse and 6-h of light during the second half of the day (aSPP-2; Fig. 1A). As for SPP, we first sought to determine the extent to which wild-type and genetic background strains entrain to aSPP regimes. We found that there was no significant reduction in the number of flies entraining to either of the asymmetric skeleton photoperiods for any genotypes (Suppl. Table 2).

We next compared the phases of the evening peak of activity under our two aSPP conditions to those seen under LD12:12 and found that, for wildtype *CS* flies, there was a significant advance in the timing of evening activity when the long-duration of light was delivered around dawn (aSSP-1; Figs. 1A and 1B). Though the evening peak appeared to be delayed when long duration light was delivered around dusk, the differences in the median phases of individual flies did not reach significance (aSPP-2; Figs. 1A and 1B). Similar effects of long-duration light around dawn were observed for *w^1118^* and *yw* flies (aSPP-1; Fig. 1) and the evening peak was significantly delayed in these strains when long duration light was delivered around dusk (aSSP-2; Fig. 1). The difference in phases of evening peak of *yw* flies under LD12:12 and aSPP-2, although significant was only ~15-min and is therefore barely apparent in the profiles (Fig. 1).

Taken together, these results reveal that long durations of light at the end of the day are sufficient for flies to achieve a waveform similar to that displayed under LD12:12. Therefore, entrained waveforms of locomotor activity under standard laboratory LD conditions are strongly shaped by time-of-day dependent continuous effects of light.

### Wildtype and background strains have longer free-running periods under constant blue- and UV-blocked light

While the number of discrete transitions between light and dark are the same across the different skeleton photoperiod regimes, the timings of these transitions are different (see Fig. 1A). This may therefore appear as a confound in our experimental design. However, instantaneous effects of light on the clock around mid-day (which is where our transitions occur) are almost non-existent [2,18,35,36]. Thus, our results described above are consistent with the speed of internal clocks being modulated by long durations, and continuous effects, of light. The most direct method available for measuring such light effects is the demonstration that free-running periods of a circadian rhythm differ between constant light and constant dark conditions. Given the fact that the presence of the blue light circadian photoreceptor CRY renders the *Drosophila* clock arrhythmic under white LL, we assayed locomotor activity in flies under various light conditions with differing degrees of overlap with the CRY action spectrum (Fig. 2A). We assayed *CS* flies under constant green, orange, yellow, blue- and UV-blocked and red lights and found that flies were arrhythmic under green and yellow lights (see example actograms in Fig. 2B). While many flies were arrhythmic under orange light, some flies displayed complex rhythms (Fig. 2B) in which spontaneous changes in free-running period are apparent. Under constant red light, flies were rhythmic and had periodicities similar to those measured under constant darkness (a little less than 24-h; Fig. 2B). In contrast, we found that when the blue- and UV-components of white light are blocked, flies are robustly rhythmic and are characterized by periodicities longer than those measured under constant darkness (Fig. 2B; Suppl. Fig. 1A; Suppl. Table 1). Thus, the presence and speed of endogenous rhythms appear to depend on the extent of overlap between the spectrum of constant light used and the action spectrum of CRY. There is considerably larger overlap in case of green and yellow light (Fig. 2A), conditions under which arrhythmicity was observed. In case of orange light, there is relatively little overlap, and this was associated with of complex rhythms in a subset of flies. The complete lack of overlap in case of red light does not seem to affect the presence or speed of the clock. However, the blue- and UV-blocked light, which has minimal but non-zero overlap with the CRY absorption spectrum, supports robust but slower rhythms. Taken together, these results suggest that CRY may play a role in the regulation of clock speed during prolonged illumination when light conditions do not drive the CRY mediated loss of circadian timekeeping.

Before examining the role of CRY on rhythm speed, we performed dose response analyses to examine effects of light intensity on the free-running period of the clock driving rhythms in locomotor activity. We used *CS*, *w^1118^* and *yw* under three intensities of constant blue- and UV-blocked light. Fitting the logistic model (see Methods) indicated that there was significant light intensity dependent change in speed of the locomotor activity rhythm in all three fly strains (Fig. 3A). However, the extent of change in rhythm speed depended on the strain used (Figs. 3A-C). While *CS* flies displayed a period lengthening of ~1-h, the *w^1118^* and *yw* strains displayed lengthening of ~3 to ~3.5-h (Figs. 3A-C). We also examined the strength of these rhythms under constant light and constant darkness (see Methods) and found that *w^1118^* and *yw* flies had significantly lower rhythmic strengths under LL conditions (Suppl. Fig. 1A; Suppl. Table 1). *w^1118^* and *yw* strains lack normal screening pigment in the eyes due to a loss of function mutation in the *white* gene. Our results therefore suggest that external photoreceptors may play a role in light dependent changes in rhythm strength and speed, owing to the *white* gene’s role in photoreception e.g., [37].

### Continuous effects of light are dependent on both external photoreceptors and *cryptochrome*

The *Drosophila* circadian system has two major photoreceptive systems, one mediated by the deep brain, cell autonomous blue-light photoreceptor *cryptochrome (cry)*, and the other consisting of external photoreceptors, i.e, the eyes, ocelli and the HB eyelets [38]. Previous work by others has established that external photoreceptors and CRY cooperate to entrain the fly’s circadian clock to 12:12LD cycles, e.g., [39]. Our results also suggest that both CRY and external photoreceptors likely contribute to light dependent changes in endogenous period. We therefore expanded our analyses to fly strains that have mutations in these light input pathways. The loss-of-function *cry^OUT^* mutation lacks functional *cry* [20] and the *glass^[60j]^* mutant lacks all external photoreceptors due to the loss of a transcription factor required for their development [13]. Previous work has shown that *cry/glass* double mutants are circadian blind to environmental light, suggesting that, taken together, *cry* and external photoreceptors account for most if not all light input to the circadian clock [13]. Further, studies have also shown that *cry* may have the ability to integrate dim light information over long durations and that *cry* mutants continue to display weak resetting responses to light [35,40].

Therefore, to further examine the extent to which external photoreceptors and CRY contribute to the parametric effects of light, we examined the effects of constant light on mutants lacking normal photoreception/transduction. We found that loss-of-function *cry^OUT^* mutants do not increase their free-running period under continuous light; rather, they showed a significant *reduction* in free-running period compared to their values under constant darkness, although this difference was quite small. This indicates that CRY is sensitive to long wavelengths light under prolonged illumination, and that it contributes to parametric light effects under a broader range of wavelengths than expected from its action spectra [24] (Figs. 3B and C; Suppl. Table 1). Remarkably, loss-of-function *cryptochrome* mutants show twice as much advance in the phase of their evening peaks of activity under SPP compared to their background controls (compare Fig. 1 and Suppl. Fig. 2). This is also consistent with the idea that *cryptochrome* tends to lengthen period under long durations of light.

We also found that a large proportion of *glass^[60j]^* mutants were arrhythmic under constant light, but that the free-running period of rhythmic flies was longer under constant light by ~1.75-h (Figs. 3B and 3C; Suppl. Table 1). In addition, we found that all *norpA* mutants, which lack the phopholipase-C signaling that mediates rhodopsin-based phototransduction, were rhythmic and their free-running periods were longer under constant light by ~3.5-h (Figs. 3A and 3C; Suppl. Table 1). Thus, in contrast to *cry^OUT^* mutants, which showed a very small reduction in period under constant light, flies lacking external photoreceptors were characterized by large parametric increases in period.

When we compared the rhythmic strength of these mutants under constant light and darkness, we found that while *cry^OUT^* displayed comparable rhythmic strength under the two regimes (Suppl. Fig. 1B; Suppl. Table 1), both *glass^[60j]^* and *norpA* both displayed significantly weaker rhythms under constant light (Suppl. Fig. 1B; Table 1). Thus, the loss of *cry* function drastically decreases the parametric effects of light, while the loss of external photoreceptor function increases them and weakens endogenous rhythms under constant light conditions.

These results suggest that parametric effects of light modulate clock speed in *Drosophila* and suggest that these are likely to contribute to light entrainment. Surprisingly, *cryptochrome* appears to be required for such lengthening of free-running periods, even under light conditions lacking blue and UV light. Furthermore, the increased extent of period lengthening by constant light in *norpA* mutants compared to wildtype *CS* flies suggests that phospholipase-C dependent phototransduction normally acts in opposition to *cry* with regard to the parametric effects of light, a result that supports previous work supporting antagonistic roles for *cry* and external photoreceptors in the adjustment of the evening peak of activity under step function LD cycles [39] and suggesting that parametric light effects make significant contributions to the timing of the entrained evening peak of activity.

### Ramped blue- and UV-blocked light cycles have significant experimental utility for understanding entrainment

Although our skeleton photoperiod experiments revealed that entrained waveforms of locomotor activity under LD12:12 are significantly shaped by parametric effects of light, the large startle responses to the discrete transitions between darkness and light which can be seen clearly in the activity profiles (Fig. 1), may contribute to the daily timing of activity in ways that may obscure the effects of light on the clock. For example, arousal signals caused by the masking effects of light likely contribute significantly to the timing of major peaks of activity. Thus, such startle responses pose a challenge to understanding the true contribution of the entrained circadian system to these waveforms. Moreover, waveforms of loss-of-function clock and light input mutants look remarkably normal under step function LD cycles in that they reliably produce two daily peaks of activity at dawn and dusk [13,41]. To characterize the timing of entrained circadian outputs most accurately there is therefore a need for a light entrainment paradigm to which wildtype flies readily entrain but that do not induce strong diurnal rhythms in loss-of-function clock mutants. Furthermore, light regimes that better reveal entrainment deficits in flies lacking normal light input pathways would be of great utility to the field. Experiments conducted under such light regimes, although structurally very different than widely used step-function LD cycles, could reveal neuronal and molecular underpinnings of circadian entrainment that step-function LD cycles will not.

To this end, we asked if ramping light cycles in which light intensity gradually changes from a low but non-zero value to a high value and then returns to the low value over 24-h, might provide novel experimental utility. We argue that such a regime would be useful because (i) there are no discrete jumps between states of darkness and light and therefore no cause for large startle responses, (ii) ramping of the intensity represents a much more challenging light condition for entrainment than step function LD cycles, as it requires the circadian system to constantly monitor a slowly changing environment, and (iii) there is no opportunity under such a light cycle for the discrete phase-shifts of the clock. If so, ramping light cycles would offer a unique regime of entrainment biased toward the parametric effects of light, making it a useful complement to the widely employed LD12:12 cycle. Light ramps might, therefore, make possible new approaches to understanding the mechanisms underlying this important form of circadian entrainment.

To examine the potential utility of ramping light cycles, we placed our wild-type, genetic background, and several mutant strains under ramping light cycles that were filtered for UV and blue wavelengths (see Methods) and characterized individual flies as either (i) entrained, (ii) displaying relative coordination (transient or unstable entrainment), (iii) free-running, or (iv) arrhythmic (Fig. 4). We found that while close to 80% of wildtype *CS* flies entrained to yellow ramps, only ~50% of *w^1118^* and *yw* flies entrain. Interestingly, ~80% of *cry^OUT^* and ~97% of *norpA* mutants displayed free-running rhythms under these conditions (Fig. 4; Suppl. Table 3) indicating the presence of a strong circadian oscillation that had failed to entrain to the light cycle. In the case of *glass^[60j]^* mutants, ~50% of the flies entrained and ~40% of flies are arrhythmic under these ramping light cycles (Fig. 4; Suppl. Table 3).

**Figure 4:**
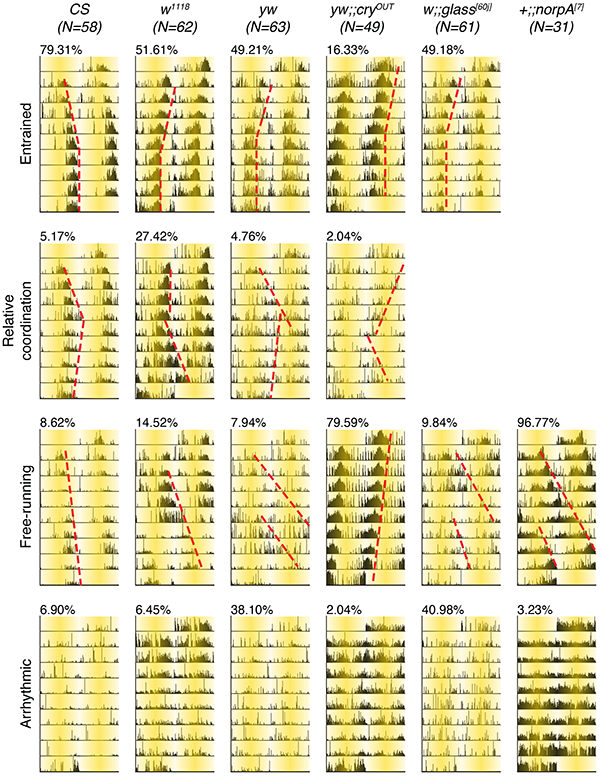
Example actograms from all six genotypes illustrating entrainment, relative coordination, free-running and/or arrhythmicity. The red dashed lines are drawn to trace the offset of locomotor activity in the top three rows. The percentage values on top of each actogram are the percentage of all flies showing that behavior. The shading on the actograms represents the ramping up and down of light, starting at Zeitgeber Time-00.

We examined the locomotor activity profiles of entrained wildtype *CS*, *w^1118^* and *yw* flies under these conditions, and found one major peak of activity that was approximately centered around the second half of the declining phase of light intensity (Fig. 5A). There is also a smaller peak of activity especially visible in *w^1118^* flies, which commences at the beginning of the inclining phase of light intensity (Fig. 5A). Interestingly, while the median phase of the dominant peak of activity for *CS* and *w^1118^* flies was ~ZT19 (with ZT00 here defined as the start of the inclining light phase), the median phase of the dominant peak in *yw* flies was ~ZT17 (Fig. 7B), a difference of ~2-h in the phase of entrained activity peaks that is not apparent under standard LD12:12 entrainment conditions (e.g., Fig. 3C and D). This surprisingly large difference between strains under light ramps but not under LD12:12 lends strong support to our notion that such light ramp cycles will allow the experimentalist to detect the effects of genetic and neuronal manipulations on entrainment that would not be apparent using standard LD12:12 cycles, thanks, most likely, to the absence of startle/arousal mediated effects on the timing of behavior and to the biasing of entrainment toward parametric effects of light.

**Figure 5:**
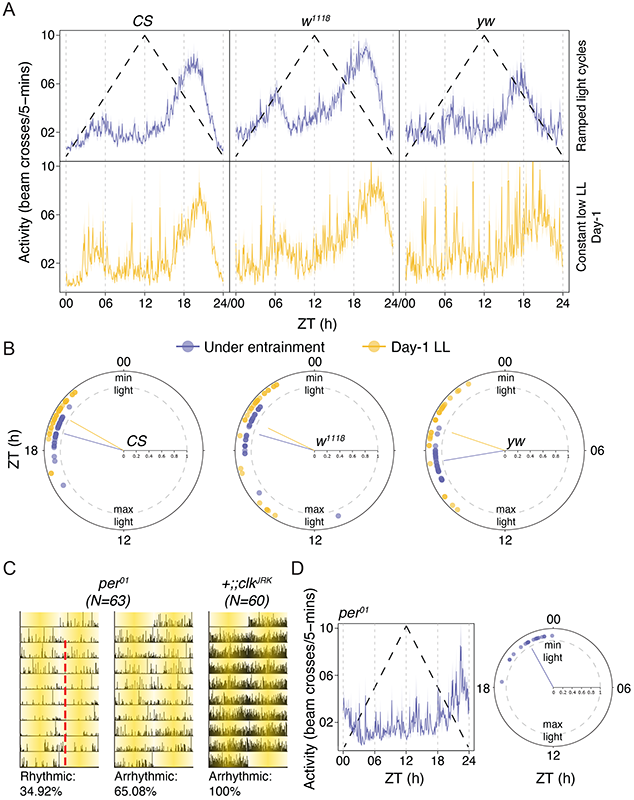
Entrainment to ramped blue- and UV-blocked light cycles. (A) Mean (±SEM) locomotor activity profiles of entrained *CS* (left), *w^1118^* (middle) and *yw* (right) flies under ramped light cycles (top) and on the first day under constant low intensity light (bottom). (B) Polar plots showing the phases of the predominant peak of locomotor activity for the three genotypes (*CS* – left; *w^1118^* – middle; *yw* – right) under entrainment and on the first day post entrainment under constant conditions. Phase control was statistically examined using V-tests (see Methods). *CS*: *p* = 7.67e-17; *w^1118^*: *p* = 1.45e-10; *yw*: *p* = 7.38e-10. (C) Representative actograms showing rhythmic and arrhythmic flies under ramped yellow cycles for loss-of-function clock mutants. The red dashed line indicates the offset of locomotor activity. The shading on the actograms represents the ramping up and down of light, starting at Zeitgeber Time (ZT) 00. (D) Mean (±SEM) locomotor activity profile (left) and polar plot depicting phases of the peak of activity of *per^01^* flies under ramped light cycles. The black dashed line in the top row of panel (A) and (D-left) is a schematic representation of light ramping from a lower minimum value (~10 *μW/cm^2^*) at ZT00 to the maximum value (40-50 *μW/cm^2^*) at ZT12 and back to the minimum value by ZT24. In all the polar plots individual dots represent a single fly and the line from the center points to the mean phase across all flies. The distance of these lines from the center is an estimate of the across-fly variation in phase. Values close to 1 (gray dashed circle) have lower variances.

A critical property of entrainment is phase control, which means that once allowed to free-run following entrainment to a cyclic environment, organisms must show similar phases of rhythm on the first day of constant conditions as it did during stable entrainment [2]. In the absence of phase control, one cannot conclude that the organism was really entrained during their exposure to any particular environmental cycle (add same references as above in methods). We estimated phase control in the wild-type and background control strains and found that phases of the dominant peak of *CS* flies on the first day under constant conditions is significantly clustered around the median phase of the same flies under entrainment (Fig. 5B). We found similar significant clustering in case of the *w^1118^* (Fig. 5B) and *yw* strains (Fig. 5B), thereby indicating that these flies had truly entrained to the previous light ramps rather than being driven directly by changes in environmental light.

To further establish the extent to which our ramped light cycles provide experimental conditions that reduce the masking effects of light on behavior, we examined loss-of-function clock mutants under these conditions. We found that only of ~35% of *per^01^* loss-of-function mutants displayed some form of rhythmicity, with most flies displaying arrhythmicity (~65%; Fig. 5C; Suppl. Table 4). We note that the residual rhythmicity was much weaker than what is observed under LD12:12, a condition in which the majority of *per^01^* mutant display rhythms. In the *per^01^* flies that were rhythmic, there was a single peak of activity very close to ZT00 or the time of the day when light intensity was at a minimum (Fig. 5D). We quantified their phases and found that the median phase of activity was ~ZT22.29. We think that such residual rhythmicity in *per^01^* flies is some form of bright light avoidance behavior/masking. We found that this phase of peak is significantly delayed compared to the phase of their background control *CS* flies (Wilcoxon’s test: *W* = 36, *p* = 9.3e-09), which is not typically the case under standard LD12:12 conditions, wherein evening peaks coincide with dusk. For *clk^JRK^* mutants, which bear a loss-of-function mutation of the *clock* gene, all flies were arrhythmic under light ramps (Fig. 5C), which is in stark contrast to the relatively nocturnal activity rhythms displayed by this mutant under LD12:12 [42]. These results suggest that light ramp cycles will offer significantly increased experimental utility for the examination of light entrainment in *Drosophila*.

## Discussion

Real world entrainment is mediated by a combined contribution of parametric and non-parametric effects of light. Parametric effects of light have been modeled in the past and have successfully explained entrainment under various photoperiodic conditions in *Neurospora*, *Drosophila* and mammals [8–10], unlike the non-parametric model which has limited ability to predict entrainment under different photoperiods [5,33]. In fact, the non-parametric model alone also has limited utility in predicting entrainment in humans (see [8] and references therein, and [43]). Therefore, a holistic study of entrainment requires the understanding of both parametric and non-parametric processes, and our study is a significant first step in that direction for *Drosophila*.

### Antagonistic effects of CRY and norpA in parametric light entrainment

Our results show that wildtype flies have longer free-running periods under constant light. The amount of this lengthening appears to be strongly dependent on both *cry* and *norpA* mediated phototransduction. Not only does the loss of CRY prevent constant light from lengthening free-running period, but it also appears to modestly reduce it. On the other hand, the loss of *norpA* function results in larger LL-induced increases in period compared to wildtype controls (see Fig. 3 and Suppl. Table 1). These clearly indicate an antagonistic interaction between *cryptochrome* and *norpA* under constant light. The idea of such an antagonism is also supported by studies in flies showing opposite effects of *cry* and visual photoreception on the evening peak of locomotor activity and their effects on clock oscillations under long days [39].

Such an antagonistic relationship can be hypothesized to be of adaptive value in the context of parametric entrainment, wherein organisms must modulate the speed of their clocks in a time-of-day dependent manner. Such antagonism can explain the entrainment of locomotor rhythms to skeleton photoperiods. *CS* flies displayed a significantly advanced phase of evening peak activity under the symmetrical skeleton photoperiod (SPP) relative to LD12:12, consistent with the idea that the absence of a full 12-h photoperiod leads to a faster running clock, thereby advancing evening peak phase under SPP. Similar results in the *w^1118^* and *yw* background strains lend further support to this idea.

The significant evening peak advances displayed by wild-type and background strains under SPP indicates that the two brief light pulses do not comprise the minimum required light schedule to recapitulate LD12:12 behavior. We therefore extended our analyses to asymmetric SPPs to ask if longer durations of light are required at specific times of day for activity to be timed in manner similar to that observed under standard LD12:12. In this way we tested the prediction made by the parametric model of entrainment that long durations of light contiguous with dawn would have a net accelerating effect on the clock while long durations of light contiguous with dusk would have a net decelerating effect on the clock. Indeed, this is exactly what we found: wild-type and genetic background flies had significantly advanced phase under long dawn skeletons and a tendency towards increased similarity to LD12:12 under long dusk skeletons. To the best of our knowledge, this is the first study to demonstrate that long durations of light at specific times of the day are required for entrainment phases like those observed under LD12:12 in the locomotor activity rhythm of flies. Similar experiments have been performed to examine the light requirement schedules of the eclosion rhythm for *D. melanogaster* flies that were artificially selected for morning and evening adult emergence [32]. This study found that for genotypes that had shorter than 24-h periods under constant darkness and a larger delay/advance ratio of their phase response curves, regimes that had long durations of light contiguous with dusk comprised the minimum light requirement schedule to mirror behavior under LD12:12 [32], providing further support for time-of-day specific parametric light effects.

### Cryptochrome mediates the parametric effects of light in D. melanogaster

The most direct effect of light on the molecular circadian clock in *D. melanogaster* is through the deep brain blue light photoreceptor CRY, which has long been thought to mediate the non-parametric effects of light the circadian clock by driving the rapid degradation of TIM upon exposure to light, thereby causing phase dependent advances or delays in the molecular circadian oscillator (e.g., [11]). However, the construction of light phase response curves (plots of time-of-day dependent phase-shifts in response to brief pulses of light) for the *cry^baby^* mutant, revealed that parametric light effects were still present, though reduced, in the absence of CRY function [35]. The presence of CRY is also thought to explain why dipteran clocks cannot maintain their rhythms under constant light conditions: CRY action under continuous presence of light prevents sufficient build-up of TIM for the molecular clock to run [44–46]. Given that CRY has long been thought to be sensitive specifically to UV/blue wavelengths, we assumed that our blue- and UV-blocked regime would “work-around” CRY. Indeed, the strong endogenous timekeeping displayed by wild-type flies under this regime was reminiscent of the rhythms displayed by loss of function *cry* mutants under the continuous white light conditions that render normal flies arrhythmic [44]. This suggested that CRY was not engaged under blue- and UV-blocked light. However, the lack of parametric effects in *cry^OUT^* flies (Fig. 3) revealed, surprisingly, that non-UV/blue wavelengths likely do engage CRY and contribute to the parametric effects of light.

Owing to the clear absence of period lengthening under blue- and UV-blocked light in loss-of-function *cryptochrome* mutants, we hypothesize that CRY activation and CRY mediated TIM degradation may be at the heart of period lengthening in this regime. This is because the action spectrum of CRY suggests that it is likely to be weakly sensitive to wavelengths of light longer than 500nm (Fig. 2A) [24]. Furthermore, flies display clear behavioral preferences for specific wavelengths of light throughout the circadian cycle, avoiding blue light and seeking out green and red shifted wavelengths during the day [47]. We suspect that, in real world settings, flies are likely to be exposed, for significant durations, to light that is significantly more yellow/red shifted than the white light used in the laboratory. Therefore, under natural and freely behaving conditions, CRY mediated TIM degradation is expected to be significantly lower than that seen in the lab. This would allow CRY to mediate the parametric effect of light by controlling TIM turnover in a manner that depends on light intensity and duration, thereby adjusting PERIOD accumulation and nuclear entry to modulate the clock’s speed. This, we argue, is mechanistically similar to the effects of a hypomorphic *cry* mutation under bright white LL, which produces lengthened free-running periods [48]. Future work examining molecular clock cycling under a range of light conditions will be necessary to test this hypothesis.

### On the utility of light ramps to study entrainment

The vastly simpler nervous system and highly conserved molecular clock in *Drosophila* has made it a superlative model system to understand mechanisms of entrainment that are relevant to entrainment in animals generally. However, one aspect of light entrainment in flies has limited its relevance to light resetting in mammals: whereas mammals including humans require several cycles to entrain to a shifted (i.e., jet lagged) LD cycle (reviewed in [2]), flies adjust to even large shifts in 12:12 LD cycles almost instantaneously upon encountering a shifted light transition (reviewed in [38]), and even flies lacking major components of light input pathway re-entrain fairly rapidly (reviewed in [38]). For these reasons, new experimental light regimes are necessary if *Drosophila* is to be used to address entrainment in a manner with the greatest possible relevance to animals generally.

We wonder if this apparent instantaneous re-entrainment to shifted LD cycles is the result of the large startle responses produced by the light/dark cycles under which flies are typically studied. Standard step-function LD cycles (and our various skeleton photoperiods – Fig. 1 and Suppl. Fig. 2) promote significant startle responses at light transitions, and these, rather than light’s effect on the clock, may be driving the rapid re-entrainment seen in flies. We propose the blue- and UV-blocked ramping light cycles as one such regime, as they fail to produce the large startles associated with step-function LD cycles and require multiple cycles for entrainment (i.e., they produce transients; Fig. 4). Furthermore, unlike standard LD12:12 conditions, blue and UV-blocked light ramps reveal significant entrainment defects in flies lacking light input pathways and fail to drive strong daily rhythms in loss-of-function clock mutants. This regime, therefore, allows us to explore aspects of entrainment in flies that are highly relevant but difficult to study using step-function LD cycles.

## Conclusion

In summary, we have demonstrated that parametric effects have a strong influence on entrainment under the long-employed step function LD12:12 and that, as predicted by parametric models of entrainment, there are clear time-of-day dependent parametric effects of light. We have also established light regimes that demonstrate significant effects of continuous light on the *Drosophila* circadian clock and reveal an unexpected role for CRY in the mediation of these effects, at wavelengths beyond those associated with its established role as a blue light circadian photoreceptor. Interestingly, the apparent antagonism of CRY and phototransduction from external photoreceptors explains the patterns of entrainment displayed under various skeleton photoperiod regimes. Finally, our work establishes experimental light conditions that will offer great utility for future research aimed at understanding the parametric effects of light on the clock and the ways in which specific genes and neural pathways contribute to circadian light entrainment.

## Acknowledgments

This work was supported by a grant from the National Institute of Neurological Disorders and Stroke (R01NS077933) and start-up funds provided by the State of New York. We would like to thank Hannah Pettibone, Matthew Ciolkowski and Lukasz Widziszewsky for technical support and Rob Veline and Budha Chowdhury for comments on the manuscript.

## Supplementary Figure and Table Legends

**Suppl. Table 1**: Details of experiments performed to assay free-running locomotor activity rhythms in various genotypes of flies under constant darkness and constant light (also see Fig. 3).

**Suppl. Table 2**: Percentage of flies entraining to each of the four environmental conditions and the total number of flies based on which these percentages were computed. There was no statistically significant reduction in the percentage of entrainment for any of these strains under any photoperiodic regime. Statistical comparisons were made using the Fisher Exact Test.

**Suppl. Table 3**: Percentage of flies that belong to one of the four entrainment/rhythm categories under ramped blue- and UV-blocked light cycles, along with the total number of flies based on which these percentages were calculated.

**Suppl. Table 4**: Percentage of flies that are rhythmic and arrhythmic for loss-of-function clock mutants under ramped blue- and UV-blocked light cycles, along with the total number of flies based on which these percentages were calculated.

**Suppl. Figure 1**: Power of circadian locomotor rhythms under constant darkness (DD) and constant light (LL) conditions for background controls and photoreceptor mutants. (A) Power of the rhythm for *CS* (left), *w^1118^* (middle) and *yw* (right). Statistical comparisons are based on Wilcoxon’s tests (*CS*: *W* = 3749, *p* = 0.0071; *w^1118^*: *W* = 7579, *p* = 1.525e-12; *yw*: *W* = 3196, *p* = 4.799e-12). Values of power are pooled from at least two independent runs (see Suppl. Table 1). * < 0.05, ** < 0.01, *** < 0.001, and NS (Not Significant). (B) Power of the rhythm for three photoreceptor mutants. Statistical comparisons are based on Wilcoxon’s tests (*yw;;cry^OUT^*: *W* = 5289, *p* = 0.76; *w;;glass^[60j]^*: *W* = 1315, *p* < 0.001; *+;;norpA^[7]^*: *W* = 2365, *p* < 0.001). Values of rhythm power are pooled from at least two independent runs (see Suppl. Table 1). * < 0.05, ** < 0.01, *** < 0.001, and NS (Not Significant).

**Suppl. Figure 2**: Entrainment to white light skeleton photoperiods. (A) Mean (±SEM) locomotor activity profiles of *yw;;cry^OUT^* flies under symmetric skeleton photoperiods (left) and two asymmetric skeleton photoperiods (middle and right). The photoperiodic regimes are shown on top of the profile plots. Black shaded regions indicate darkness and white regions indicate light phases (400-500 lux) of the 24-h cycle. Note that the profiles under LD12:12 are replotted across panels to facilitate pair-wise comparisons. (B) Phases of the evening peak of activity across the different photoperiodic conditions are shown. Boxplots that share the same letter are not statistically significantly different from each other. Statistical comparison of phases across all four photoperiodic conditions were done using Kruskal-Wallis tests. 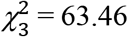 = 63.46, *p* = 1.07e-13. Profiles and phases of evening peak of activity are pooled from two independent replicate runs for each genotype.

