## Supplemental Data and Legends for "Parametric effects of light acting via multiple photoreceptors contribute to circadian entrainment in *Drosophila melanogaster*"

*Running title: Parametric entrainment in Drosophila*

**Suppl. Table 1:** Details of experiments performed to assay free-running locomotor activity rhythms in various genotypes of flies under constant darkness and constant light (also see Fig. 3).

| | ***N*** | | ***% Rhythmic*** | | ***Period (h)*** | | ***Rhythm power*** | | ***Number of independent runs*** | |
|---|---|---|---|---|---|---|---|---|---|---|
| | ***DD*** | ***LL*** | ***DD*** | ***LL*** | ***DD*** | ***LL*** | ***DD*** | ***LL*** | ***DD*** | ***LL*** |
| *CS* | 120 | 128 | 80.46 | 86.32 | 23.42 | 24.67 | 229.12 | 306.26 | 4 | 4 |
| $w^{1118}$ | 96 | 128 | 97.92 | 81.25 | 23.33 | 26.89 | 972.90 | 535.07 | 3 | 4 |
| *yw* | 113 | 126 | 96.46 | 25.47 | 24.08 | 25.17 | 270.76 | 57.16 | 4 | 4 |
| $yw;;cry^{OUT}$ | 100 | 119 | 96.77 | 100.00 | 23.42 | 23.00 | 718.05 | 729.37 | 4 | 4 |
| $w;;glass^{[60j]}$ | 119 | 128 | 81.04 | 18.15 | 24.54 | 26.39 | 171.05 | 23.40 | 4 | 4 |
| $norpA^{[7]}$ | 62 | 62 | 98.08 | 100.00 | 24.67 | 28.25 | 884.40 | 400.73 | 2 | 2 |

**Suppl. Table 2:** Percentage of flies entraining to each of the four environmental conditions and the total number of flies based on which these percentages were computed. There was no statistically significant reduction in the percentage of entrainment for any of these strains under any photoperiodic regime. Statistical comparisons were made using the Fisher Exact Test.

| | ***% Entrained (N)*** | | | |
|---|---|---|---|---|
| | ***LD12:12*** | ***SPP*** | ***aSPP-1*** | ***aSPP-2*** |
| *CS* | 100 (61) | 100 (62) | 100 (62) | 100 (64) |
| $w^{1118}$ | 100 (60) | 100 (63) | 100 (62) | 100 (58) |
| *yw* | 100 (62) | 98.38 (62) | 100 (64) | 100 (62) |

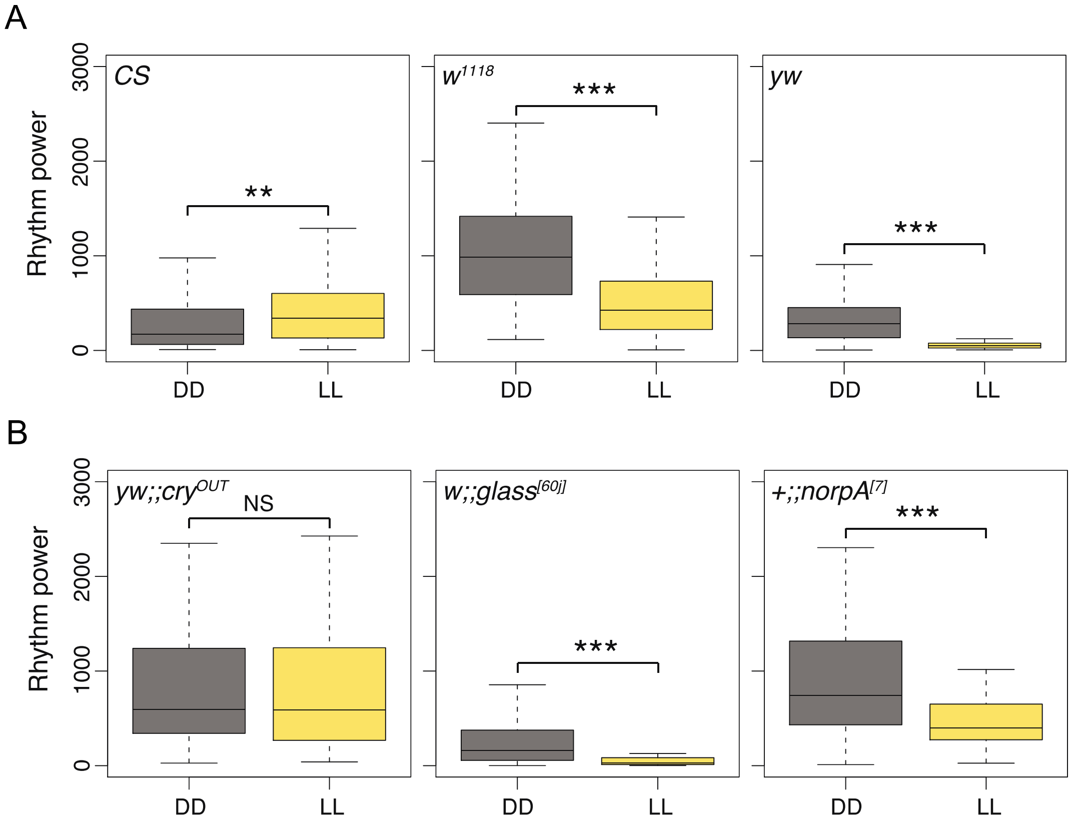


**Suppl. Figure 1:** Power of circadian locomotor rhythms under constant darkness (DD) and constant light (LL) conditions for background controls and photoreceptor mutants. (A) Power of the rhythm for *CS* (left), $w^{1118}$ (middle) and *yw* (right). Statistical comparisons are based on Wilcoxon's tests (*CS*: $W = 3749$, $p = 0.0071$; $w^{1118}$: $W = 7579$, $p = 1.525e\text{-}12$; *yw*: $W = 3196$, $p = 4.799e\text{-}12$). Values of power are pooled from at least two independent runs (see Suppl. Table 1). * < 0.05, ** < 0.01, *** < 0.001, and NS (Not Significant). (B) Power of the rhythm for three photoreceptor mutants. Statistical comparisons are based on Wilcoxon's tests ($yw;;cry^{OUT}$: $W = 5289$, $p = 0.76$; $w;;glass^{[60j]}$: $W = 1315$, $p < 0.001$; $+;;norpA^{[7]}$: $W = 2365$, $p < 0.001$). Values of rhythm power are pooled from at least two independent runs (see Suppl. Table 1). * < 0.05, ** < 0.01, *** < 0.001, and NS (Not Significant).

**Suppl. Table 3:** Percentage of flies that belong to one of the four entrainment/rhythm categories under ramped blue- and UV-blocked light cycles, along with the total number of flies based on which these percentages were calculated.

| | ***Entrained*** | ***Relative coordination*** | ***Free-running*** | ***Arrhythmic*** | ***N*** |
|---|---|---|---|---|---|
| *CS* | 79.31 | 5.17 | 8.62 | 6.90 | 58 |
| *$w^{1118}$* | 51.61 | 27.42 | 14.52 | 6.45 | 62 |
| *yw* | 49.21 | 4.76 | 7.94 | 38.10 | 63 |
| *yw;;$cry^{OUT}$* | 16.33 | 2.04 | 79.59 | 2.04 | 49 |
| *w;;$glass^{[60j]}$* | 49.18 | 0 | 9.84 | 40.98 | 61 |
| *+;;$norpA^{[7]}$* | 0 | 0 | 96.77 | 3.23 | 31 |

**Suppl. Table 4:** Percentage of flies that are rhythmic and arrhythmic for loss-of-function clock mutants under ramped blue- and UV-blocked light cycles, along with the total number of flies based on which these percentages were calculated.

| | ***Rhythmic*** | ***Arrhythmic*** | ***N*** |
|---|---|---|---|
| *+;;$per^{01}$* | 34.92 | 65.08 | 63 |
| *+;;$clk^{JRK}$* | 0 | 100 | 60 |

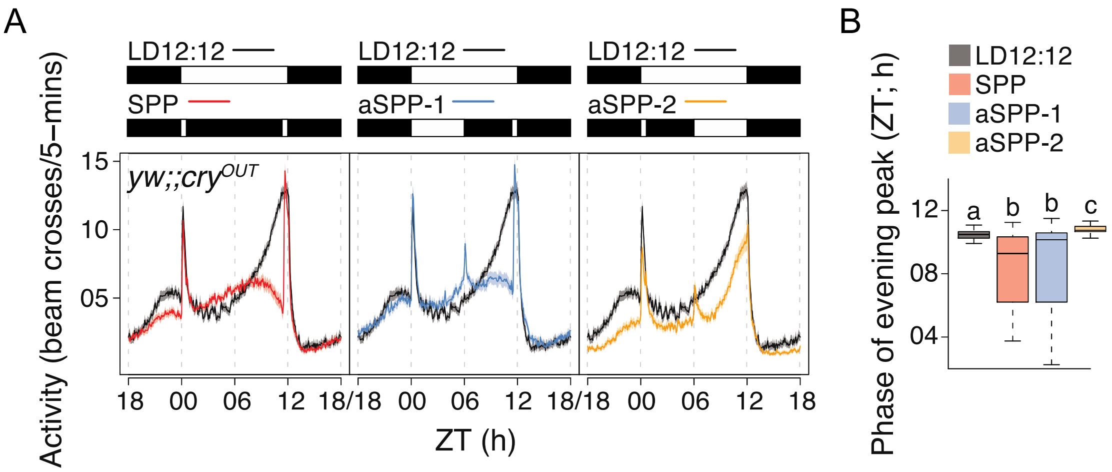


**Suppl. Figure 2:** Entrainment to white light skeleton photoperiods. (A) Mean (±SEM) locomotor activity profiles of *yw;;cry$^{OUT}$* flies under symmetric skeleton photoperiods (left) and two asymmetric skeleton photoperiods (middle and right). The photoperiodic regimes are shown on top of the profile plots. Black shaded regions indicate darkness and white regions indicate light phases (400-500 lux) of the 24-h cycle. Note that the profiles under LD12:12 are replotted across panels to facilitate pair-wise comparisons. (B) Phases of the evening peak of activity across the different photoperiodic conditions are shown. Boxplots that share the same letter are not statistically significantly different from each other. Statistical comparison of phases across all four photoperiodic conditions were done using Kruskal-Wallis tests. $\chi^2_3 = 63.46$, $p = 1.07e\text{-}13$. Profiles and phases of evening peak of activity are pooled from two independent replicate runs for each genotype.
